# Crown-of-thorns starfish in captivity experience sustained large-scale changes in gene expression

**DOI:** 10.1101/2022.07.21.501052

**Authors:** Marie Morin, Mathias Jönsson, Conan K. Wang, David J. Craik, Sandie M. Degnan, Bernard M. Degnan

## Abstract

Marine animals in the wild are often difficult to access, so that biologists have to extrapolate from the study of animals in captivity. However, the implicit assumption that physiological and cellular processes of animals in artificial environments are not significantly different from those in the wild has rarely been tested. Here we investigate the extent to which the biological state of an animal is impacted by captivity by comparing global gene expression in wild and captive crown-of-thorns starfish (COTS). We compare transcriptomes of three external tissues obtained from wild COTS with captive COTS maintained in aquaria for at least one week. On average, an astonishingly large 24% of the coding sequences in the genome are differentially expressed. Comparing transcriptomes from coelomocytes – cells in internal coelomic fluid – in wild and captive COTS, we find that 20% of the coding sequences in the genome rapidly change expression. These captive transcriptomes remained markedly different from the wild ones for more than 30 days in captivity, and showed no indication of reverting back to a wild state. Genes consistently upregulated in captivity include those involved in oxidative stress and energy metabolism, whereas genes downregulated are involved in intercellular signalling. These extensive changes in gene expression in captive COTS suggest that captivity has a profound and sustained impact on the physiology, behaviour and health of these echinoderms. The potential for such dramatic changes should be accounted for when designing studies seeking to understand wild animals.

## 1 Introduction

Animals are brought into and maintained in artificial environments for many reasons, including as models for biological, biomedical or ecological research, for species conservation, and for agriculture and aquaculture (Fischer and Romero, 2019). This is not surprising given that studying animals *in situ* is often expensive and difficult; this is especially true for subtidal marine organisms. Controlled laboratory and animal-holding facilities allow for the reduction of many variables compared to field settings, which is useful for experimental analyses and interpretation of data at almost all levels of biological enquiry (Calisi and Bentley, 2009). However, there are remarkably few empirical data addressing the implicit assumption that results from an artificial environment can reliably be extrapolated to the wild. The small number of studies that currently exist have shown that laboratory and field experiments can reveal different, and sometimes even opposite, results (Calisi and Bentley, 2009; Campbell et al., 2009; Fischer and Romero, 2019). Moreover, most studies assess changes only in specific markers of stress, such as heat shock proteins, steroid hormone concentrations or immune cell ratios (reviewed by (Calisi and Bentley, 2009; Fischer and Romero, 2019). Thus, the full extent of the impact of being in captivity is poorly understood.

Because gene expression can be influenced by environmental conditions, it is a valuable proxy for changes in an animal’s development and physiology (Oomen and Hutchings, 2017). Genome-wide transcriptome analysis using RNA sequencing (RNA-Seq) is a highly sensitive approach that provides near-comprehensive quantification of changes in gene expression under different environmental conditions (Wang et al., 2009). Although RNA-Seq has been widely applied to studying responses of diverse animals to specific environmental stressors (Lv et al., 2013; Meng et al., 2013; Milan et al., 2013; Hui et al., 2014; Gleason and Burton, 2015; Oomen and Hutchings, 2017; Zhang et al., 2017; Wang et al., 2018; Li et al., 2019; Ma et al., 2019; Li et al., 2021), it has rarely been applied in the context of captivity (Alburaki et al., 2019; Roznere et al., 2021).

Here, we use RNA-Seq to investigate the effect of captivity on global gene expression on a well-known echinoderm, the crown-of-thorns starfish (COTS, *Acanthaster planci* cf. *A. solaris*). COTS are highly fecund predators of reef-building corals living throughout the Indo-Pacific (Birkeland and Lucas, 1990; Pratchett et al., 2014), with a biology that predisposes them to population outbreaks (Moran, 1986; Birkeland and Lucas, 1990) that lead to extensive loss of live coral cover and reduced reef resilience (Yasuda et al., 2009; De’ath et al., 2012; Riegl et al., 2013), (Porter, 1972; Pratchett, 2010). Their pest status means there is an urgency to gain new insights into COTS biology to develop advanced biocontrol methods, however as a cryptic, venomous and subtidal species, they present great challenges to study in the wild. As a result, most biological and molecular studies on COTS to date have been performed in laboratory or controlled aquarium settings (Barnes et al., 1970; Brauer et al., 1970; Beach et al., 1975; Teruya et al., 2001; Petie et al., 2016a; Petie et al., 2016b; Caballes et al., 2017; Roberts et al., 2017; Smith et al., 2017; Høj et al., 2018; Smith et al., 2018; Hue et al., 2020), making it important to understand the extent to which captive studies can inform us about the wild animal condition.

To address this, we compare tissue-level gene expression profiles from COTS that were sampled directly from the wild with those maintained in captivity. We find that gene expression profiles in four different tissues in wild COTS are strikingly different from those maintained in aquaria, with over 20% of coding sequence genes in the genome being affected. Further, a detailed analysis of gene expression over one month in coelomocytes in COTS reared under different conditions of captivity (fed and unfed) reveals that the initial transition from wild to captive has the most impact, but that even one month later there is no evidence that captive COTS are reverting back to the wild state. Together, these findings reveal high levels of captivity-induced changes in gene expression across multiple different tissues.

## 2 Materials and Methods

### 2.1 Genome annotation

Using the original genome assembly (Hall et al., 2017), we use the Program to Assemble Spliced Alignments (PASA) (v2.4.1) (Haas et al., 2003) to improve the original coding sequences (CDS) models of the *Acanthaster planci* Great Barrier Reef (GBR) genome (GBR v1.0). Specifically, the Genome Mapping and Alignment Program (GMAP) (Wu and Watanabe, 2005) was used to align Trinity transcripts produced by Hall et al. (2017) to the GBR genome. Two sequential cycles of annotation loading, annotation comparison, and annotation updates were performed, as recommended in the PASA pipeline manual (http://pasapipeline.github.io/). The new GBR gene models (v1.1) contains gene isoforms and untranslated regions (UTRs). Isoforms with the longest coding sequences were identified and shorter ones were discarded, to reduce isoform redundancy. Only CDS were analyzed in this study. Gene Ontology (GO) terms and InterPro IDs were assigned to GBR v1.1 CDS using Blast2GO as part of Omicsbox (v1.4.11) using default settings (Götz et al., 2008). CDS were further annotated using the Pfam-A database (El-Gebali et al., 2019) and the Kyoto Encyclopedia of Genes and Genomes (KEGG) database using GhostKOALA searching against the genus_prokaryotes + family_eukaryotes database) (Kanehisa et al., 2016).

Previously published RNA-Seq reads of podia, spines and body wall transcriptomes from a single COTS maintained in a flow-through aquarium at the Australian Institute for Marine Science (AIMS) under winter ambient conditions for 1 to 2 weeks (Hall et al., 2017) were mapped to the GBR genome using Bowtie2 (Langmead and Salzberg, 2012). New transcript counts were generated for GBR v1.1 CDS model using HTSeq (Anders et al., 2014).

### 2.2 Collection of wild COTS

COTS were collected from Davies (18°50’S, 147°39’E) and Lynchs Reef (18°76’S, 147°63’E) on the GBR. Upon removal from the reef, COTS were transferred immediately to a vessel on the surface, where they were maintained in ambient seawater until processing. Seven and thirteen individuals were sampled in the winter (August) and summer (December), respectively. Tube feet, skin, spines and coelomocytes were isolated from each starfish within two hours of their removal from the reef. Dissected tissues were immediately transferred into RNALater, stored at 4°C for 24 hours and then at −20°C. Individual tube feet and spines were collected from the middle and base of a randomly selected arm, respectively. The skin was sampled at the proximal end at the side of the arm. 1.5 ml of coelomic fluid was collected using a syringe and a 21-gauge needle, immediately centrifuged at 2,000 x g, and the coelomocyte pellets were transferred into RNALater (Pérez-Portela and Riesgo, 2013). Papillae and pedicellaria were removed from skin samples prior to RNA extraction. The epithelial layer of spines was scrapped off and used for RNA extraction.

### 2.3 CEL-Seq2 analysis

RNA from individual COTS tissues was extracted separately using TRI Reagent (Sigma), following the manufacturer’s protocol, and assessed using a Qubit® fluorometer (Invitrogen-Life Technologies) and an Agilent 2100 Bioanalyzer (Agilent Technologies). Extracted RNA was stored at −80°C in UltraPure™ DNase/RNase-free water (ThermoFisher) until further processing. Tissues from each individual were kept separate throughout, so each tissue was a biological replicate. The CEL-Seq2 method was used to barcode individual tissue samples for RNA sequencing, with unique primers individually tagging the 3’ PolyA-tail of the transcripts in each sample (Hashimshony et al., 2016). This allowed for the subsequent pooling of samples. Wild samples were pooled by tissues and cDNA libraries were constructed following the CEL-Seq2 previously described (Hashimshony et al., 2016), The CEL-Seq2 libraries were sequenced on the Illumina Hiseq X platform at NovogeneAIT Genomics in Singapore. Raw reads were assessed for quality and adaptor contents using FastQC (Andrews, 2010), and analysed using the CEL-Seq2 pipeline (https://github.com/yanailab/CEL-Seq-pipeline) (Hashimshony et al., 2016). Reads were trimmed to 35 bp, demultiplexed and mapped to the GBR genome using Bowtie2. Transcript counts of each GBR v1.1 CDS were generated using HTSeq. Samples with < 0.5 million mapped transcripts were deemed low-quality and discarded (Sogabe et al., 2019).

### 2.4 Comparison between wild and captive transcriptomes

The original captive transcriptomes (Hall et al., 2017) were not replicated. To enable comparisons between the wild and captive datasets, we calculated a mean value of the read counts from the wild replicates and then normalised the read counts of captive and wild transcriptomes to transcripts per million (TPM) per CDS (Wagner et al., 2012). Because of the difference in sequencing depth between the RNA-Seq and CEL-Seq2 libraries, only the top two quartiles of expressed genes (i.e. transcript abundance) for each sample were analysed. CDS with a log2 fold change of ≥ ± 2 were deemed as being differentially expressed.

### 2.5 Feeding/starving experiment

Adult COTS were collected in February (summer) on the northern GBR, air transported from Cairns to Brisbane, and maintained in individual 50-litre tanks in a recirculating aquarium system in artificial seawater (Pro-reef sea salt, Tropic Marin) at The University of Queensland at 25°C under a 12/12 hours light/dark cycle. Following their arrival, COTS were starved for 5 days and randomly assigned to a feeding or starving experimental group. Four COTS were starved through the 30-day experiment and four were fed mussels to satiation twice a week. 5 ml of coelomic fluid was collected as described above from each individual at the beginning of the experiment and every 5 days for 30 days. 1.5 ml of coelomic fluid was passed through a 70 μm filter and centrifuged at 12,000 x g for 6 min at 4°C (Pérez-Portela and Riesgo, 2013). RNA was extracted from the pellet, and CEL-Seq2 sequencing and analysis were performed as described above.

For captive and wild coelomocytes samples, CDS were considered expressed if the average count per condition was ≥ 0.25. Using Wald Test statistics in the DESeq2 package (v1.28.1) (Love et al., 2014), CDS with an adjusted *p*-value < 0.05 between treatments were identified as being significantly differentially expressed. Principal component analyses (PCAs) were performed on variance-stabilise transformed (vst) counts obtained with DESeq2 to visualise variations between treatments.

### 2.6 Enrichment analyses and data visualisation

For GO and Pfam enrichment analyses, differentially expressed CDS were tested against all the genes in the COTS GBR v1.1 genome. Gene Ontology (GO) enrichment analyses of DEGs were performed using the Fisher’s exact test function available on the “clusterProfiler” package (v4.2.2) (Wu et al., 2021), using the GO annotation provided by Blast2GO. GO terms with a False Discovery Rate (FDR) adjusted *p*-value of < 0.05 were considered enriched. Pfam enrichment analysis was performed using a previously published R script, with a *p*-value cutoff of 0.05 (Chandran et al., 2009; Say and Degnan, 2020). Differentially expressed CDS were also analysed through the KEGG database using GhostKOALA (Kanehisa et al., 2016).

Data were visualised using RStudio (v4.0.2) (RStudio Team, 2020). PCAs and volcano plots were performed using ggplot2 (v3.3.3) (Wickham, 2016). Venn diagrams of GO term enrichments were made using VennDiagram (v1.6.20) (Chen and Boutros, 2011), and visualisations of common GO terms and their associated FDR-adjusted *p*-value were performed using pheatmap (v1.0.1) (Kolde, 2015). Intersecting sets of genes were visualised using UpsetR (v1.4.0) (Conway et al., 2017).

## 3 Results

### 3.1 COTS GBR v1.1 genome improvements

We used PASA on the original *Acanthaster planci* cf. *A. solaris* GBR v1.0 genome assembly (Hall et al., 2017) to update CDS. This is referred to as GBR v1.1. This update added 20,187 UTRs to 11,430 CDS models. 1,299 of the 24,747 *ab initio* gene models were merged into 623 models, resulting in 24,071 CDS and increasing genome coverage by 32% (Supplementary Table S1.1). GO analysis (Götz et al., 2008) annotated 15,944 CDS (66%) with 4,290 different GO terms. 11,396 unique Pfam domains were assigned to 18,530 CDS (77%), and 10,042 CDS (41.7%) were assigned to 3,365 KEGG Orthology (KO) groups in 401 pathways (Supplementary Tables S1.2-1.4).

New transcript counts for podia, spines, and body wall transcriptomes from captive COTS (Hall et al., 2017) were generated from GBR v1.1 gene models, revealing 37.8% reads counted; only 11.9% reads were counted with v1.0 gene models (Supplementary Table S1.5).

### 3.2 Large gene expression differences in captive and wild COTS

Seven tube feet, six spine and seven skin CEL-Seq2 transcriptomes were generated from tissues obtained from seven COTS immediately after being collected from Davies Reef (GBR) in the winter. On average, 8.5 million reads mapped to GBR v1.1, of which 50% aligned to CDS and were counted (Supplementary Table S1.6). These transcriptomes were compared with previously published transcriptomes of podia, spines, and body wall from COTS maintained in captivity in the winter (Hall et al., 2017). This comparison was performed on the top two quartiles of expressed genes (TPM-normalised). CDS were deemed differentially expressed if they exhibited greater than or equal to ±2 log2 fold change.

5,555, 6,268, and 5,449 CDS are differentially expressed in tube feet, spines, and skin respectively (Figure 1A-C), which comprise about 40% of the expressed CDS in the top two quartiles (Supplementary Table S2.1). On average, 72% of the differentially expressed CDS are upregulated in captivity. GO, Pfam and KEGG analyses of up- and downregulated CDS in each tissue reveal that upregulated genes in all three tissues in captive COTS are commonly enriched in four GO terms related to stress (Figure 1D-E; Supplementary Tables S2.2-S2.10). Other shared GO terms in upregulated genes include processes related to catabolic metabolism pathways, transcription, translation, folding, sorting and degradation, replication and repair, and thermogenesis.

**Figure 1.**
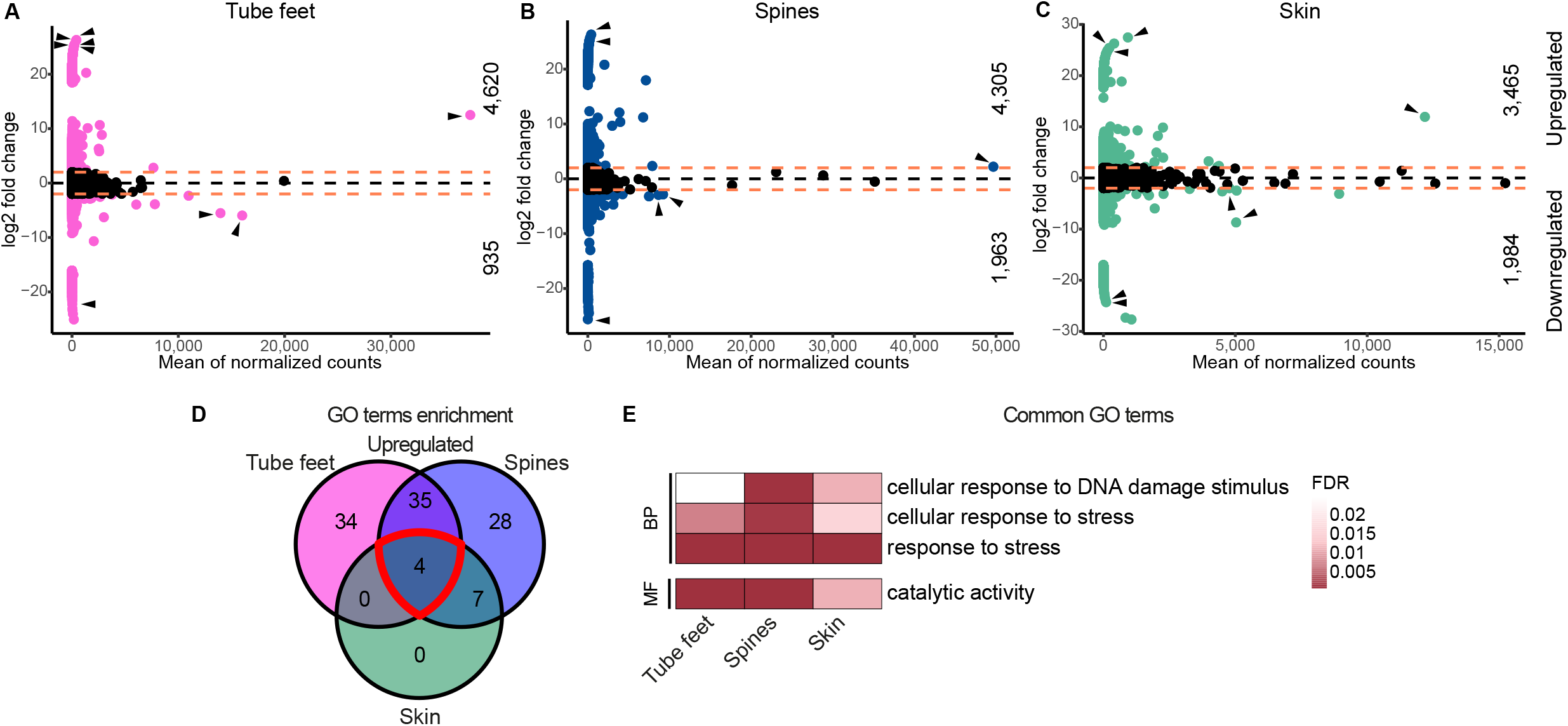
Comparison of captive and wild gene expression levels. Volcano plots of the top 2 quartiles of expressed genes in **(A)** tube feet, **(B)** spines, and **(C)** skin between captive and wild transcriptomes. Coloured dots represent differentially expressed genes with a log2 fold change ≥ ± 2 (orange dotted lines). The number of genes upregulated and downregulated in captivity are shown in the top and bottom halves, respectively. Black arrowheads point toward uncharacterised genes identified among the top five genes with highest mean and/or highest log2 fold change. (**D)** Venn diagram summarising the number of GO terms enriched in upregulated genes in captivity. Red line highlights the 4 terms common to all three tissues. (**E)** Visualisation of the 4 GO terms enriched in upregulated genes in captivity in all three tissues, and the FDR-adjusted p-value for each tissue. BP, biological processes; MF, molecular function.

Notably, genes with the highest mean expression and/or highest log2 fold change (Figure 1A-C) encode for uncharacterised proteins. Eleven of these (nine novel, two conserved) were upregulated in at least one tissue, nine (three novel, six conserved) were downregulated in at least one tissue.

### 3.3 Captivity induces quick and sustained changes in COTS coelomocytes transcriptomes

To further investigate the effect of captivity on COTS, we generated transcriptomes from cells circulating in the internal coelomic fluid – the coelomocytes – from wild and captive COTS. Coelomic cells in echinoderms and other bilaterians are involved in production and/or transport of metabolites, nutrients, and signalling and immunological factors (Shabelnikov et al., 2019). Echinoderm coelomocytes are sensitive to environmental stress and can reflect the physiological state of an individual (Branco et al., 2013; Shabelnikov et al., 2019; Hamel et al., 2021). We compared gene expression in coelomocytes of wild COTS with COTS placed in captivity and either fed or not fed over 30 days. Four COTS were fed and four were unfed. Coelomocyte transcriptomes were analysed from these eight individuals on day 0 of the experiment (after 5 days of starvation post-arrival in the aquarium), 15 and 30. Wild coelomocyte transcriptomes were derived from 13 individuals collected during the same season (summer). On average, 7.9 and 7.3 million reads mapped to GBR v1.1, of which 47% and 48% aligned to CDS and were counted for the captive and wild transcriptomes, respectively (Supplementary Table S1.6).

Regardless of the amount of time spent in captivity and feeding status, there is a general and strong impact of being held in captivity on COTS coelomocyte gene expression (Figure 2A), with 4,839 CDS being significantly differentially expressed (adjusted *p*-value < 0.05) in coelomocytes between wild and captive (20% of CDS in the genome). 69% of these are downregulated in the captive starfish (Supplementary Table S3.1), in contrast to tube feet, skin and spines where most genes are upregulated (Figure 1A-C). Two distinct groups are observed within the coelomocyte transcriptomes from wild COTS (Figure 2A), with 2,747 genes differentially expressed between the wild groups. However, the differences between wild and captive COTS appear to be greater than the differences among wild individuals.

**Figure 2.**
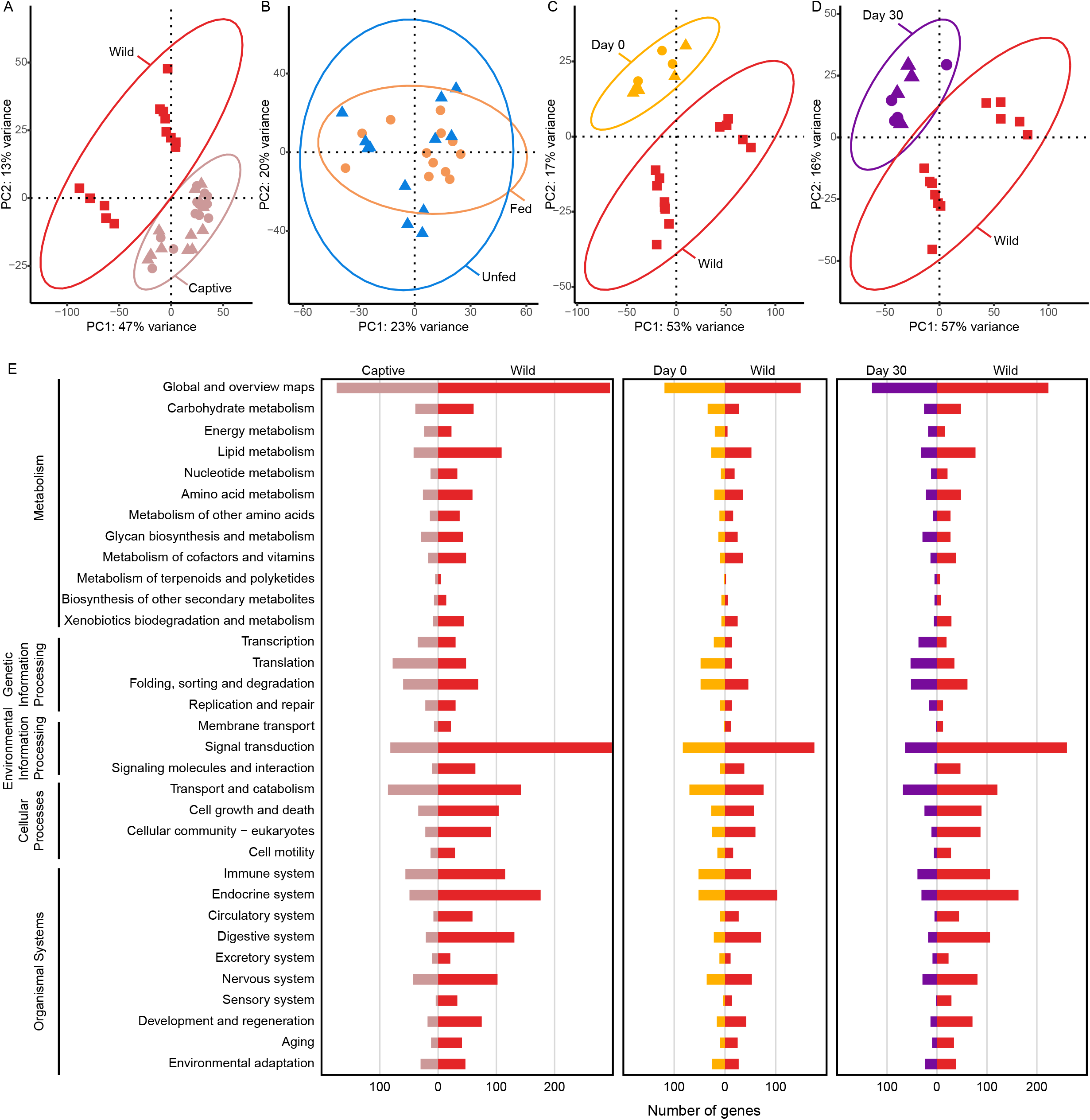
Comparison of the coelomocyte transcriptomes. **(A)** PCA of wild and captive COTS coelomocyte transcriptomes. Captive samples include all time points and feeding status. (**B)** PCA of the fed and unfed captive individuals, all time points considered. (**C)** PCA of wild and captive (experimental day 0) coelomocyte samples. Fed and unfed symbols on the plot indicate the group to which individuals were assigned after the day 0 sampling. (**D)** PCA of wild COTS samples and captive samples collected at day 30 of the experiment, both feeding status included. Dot, fed; triangle, unfed; square, wild. (**E)** KEGG analyses of differentially expressed genes in three comparisons (Captive vs Wild coelomocytes; Day 0 vs Wild coelomocytes; Day 30 vs Wild coelomocytes). Bars, number of unique genes in each category.

Only 186 genes are differentially expressed between the fed and unfed groups in captivity over the 30-day experiment, all time points included (Figure 2B). This suggests that being in captivity has more effect on the coelomocyte gene expression than the feeding status of the individuals.

GO, Pfam and KEGG analyses suggest that genes upregulated in coelomocytes of captive COTS are involved in energy metabolism, transcription and translation, whereas the downregulated genes are involved in the lipid metabolism, and endocrine, signalling, nervous and sensory systems (Figure 2E) (Supplementary tables S3.2-S3.4).

We compared both day 0 and day 30 coelomocyte transcriptomes to wild coelomocyte transcriptomes to determine whether transferring COTS from the wild into captivity has more impact than maintaining the COTS in captivity. Experimental day 0 COTS were collected from the GBR approximately 10 days earlier (5 days from initial collection to arriving in Brisbane plus 5 days of being maintained in an unfed state in aquaria). 2,956 CDS (12% of CDS in the genome) are differentially expressed between these and wild COTS coelomocytes (Figure 2C) (Supplementary Table S3.1), revealing that the time in transportation and captivity markedly affects gene expression in these cells. GO, Pfam and KEGG analyses indicate that genes upregulated in captive day 0 COTS are enriched in terms related to immunity, oxidoreduction and proteolysis, whereas genes downregulated are enriched in pathways related to cell signalling (Figure 2E) (Supplementary Tables S3.5-S3.7).

The day 30 captive coelomocyte transcriptomes have 3,883 CDS (16% of CDS in the genome) differentially expressed when compared with wild coelomocyte transcriptomes (Figure 2D) (Supplementary Table S3.1), with up- and downregulated genes enriched in terms related to DNA repair and cell signalling, respectively (Figure 2E) (Supplementary Tables S3.8-S3.10). There are 1,315 CDS differentially expressed between day 0 and 30 captive coelomocyte transcriptomes, which is more than three times less than the differences between captive and wild coelomocyte transcriptomes.

### 3.4 195 CDS are differentially expressed in all four tissues in captive COTS

68 and 127 CDS are consistently up- and downregulated in tube feet, spines, skin and coelomocytes of captive COTS compared to wild COTS (Figure 3A; Supplementary Table S4.1), potentially revealing a whole-organism response to captivity. KEGG pathway analysis reveals that genes upregulated in captivity are involved in metabolism, translation, folding, sorting and degradation, and DNA replication and repair, whereas genes downregulated are involved in signal transduction, and immune, endocrine, nervous and sensory systems (Figure 3B; Supplementary Table S4.2). Moreover, eleven conserved uncharacterized CDS, and eleven novel unannotated CDS are consistently downregulated in captivity. Six transcription factors (paired box protein Pax-6-like, nuclear receptor retinoic X receptor (RXR), zinc-finger protein GLIS3, ETS-related transcription factor Elf-1-like, photoreceptor-specific nuclear receptor, and zinc finger BED domain-containing protein 1-like) are also consistently downregulated in captivity.

**Figure 3.**
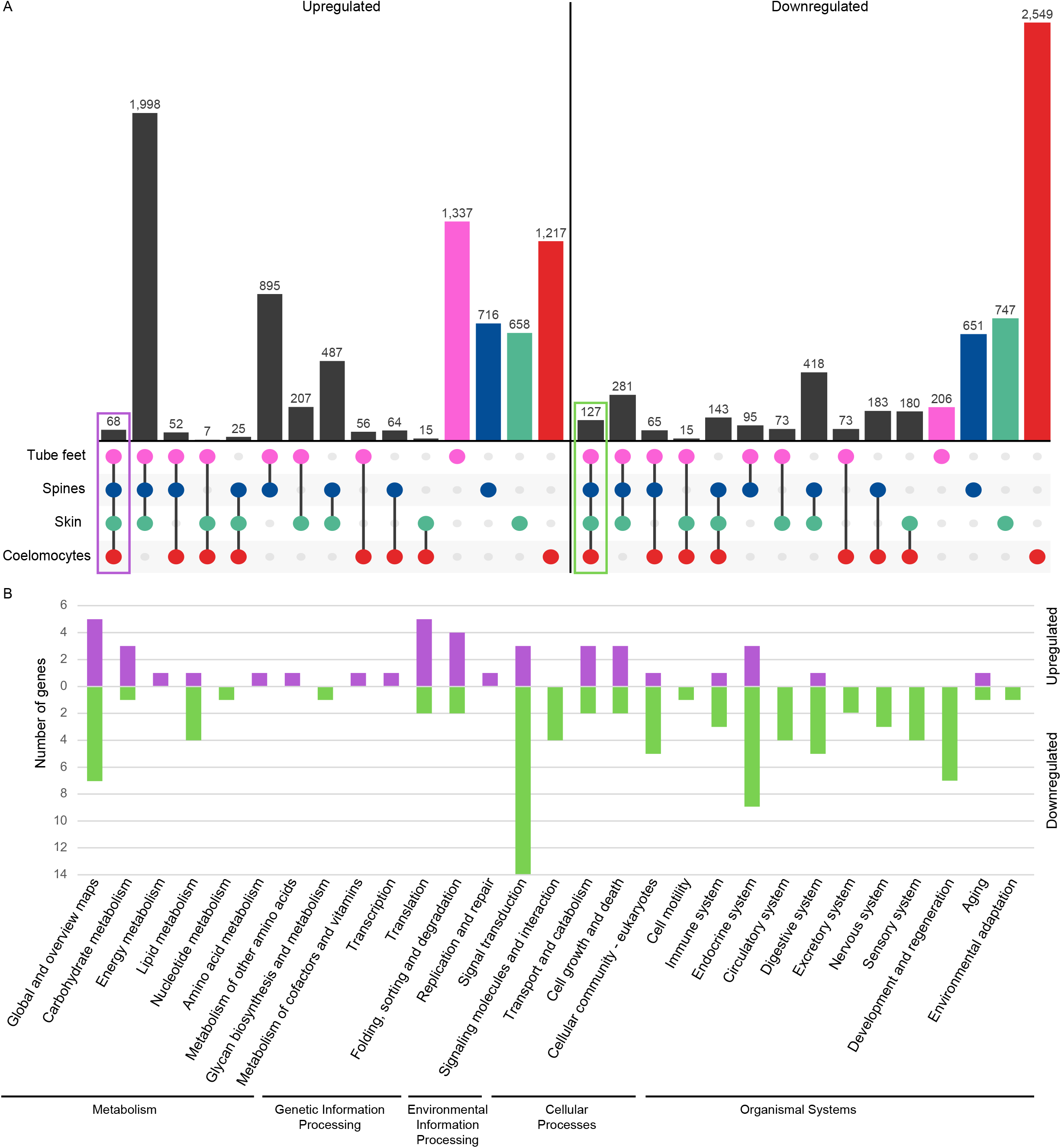
Commonality between captive transcriptomes. **(A)** Intersection plot of genes upregulated (left) and downregulated (right) in captivity in each tissue. The interaction size is the number of genes differentially expressed in one or more tissues as shown at the bottom of the graph. The purple and green boxes highlight the 68 and 127 CDS commonly up- and downregulated in captive COTS tissues, respectively. **(B)** KEGG pathways analysis of CDS up (purple)- and downregulated (green) in captive COTS tissues Bars, number of unique genes in each category.

## 4 Discussion

Although animals are regularly brought into captivity for research purposes, our understanding of the effect of laboratory environments on these organisms is limited, especially for marine invertebrates. In this study, we investigated genome-wide differences in gene expression between captive and wild COTS. To enable the highest sensitivity of our analysis, we first improved the annotation of CDS genes in the genome, which markedly increased the proportion of RNA-Seq and CEL-Seq2 reads mapping to CDS.

### 4.1 Wild and captive COTS have strikingly different gene expression profiles

When comparing tube feet, spine and skin transcriptomes from captive and wild COTS, a very high 24% of the CDS in the genome are differentially expressed on average. This comparison used captive transcriptomes generated in a previous study that were not replicated and were sequenced to a different depth (Hall et al. 2017) from the replicated transcriptomes from wild COTS generated in this study. To account for these differences, we first normalized gene expressed to TPM in all transcriptomes and then only compared relative expression level of the most highly expressed CDS (top two quartiles). Importantly, we followed this initial analysis with an experiment using a different tissue (coelomocytes) that was obtained from multiple wild and captive individuals. This allowed statistical analysis of gene expression using DESeq (Love et al., 2014). In the latter analysis (coelomocytes), 20% of the COTS CDS are differentially expressed between wild and captive COTS, similar to that observed in the tissues used in the former analysis (tube feet, spine and skin).

These large changes in overall gene expression in COTS in captivity appear to be regardless of (i) season, (ii) mode of captivity – COTS were maintained either in a state-of-the-art flow through aquarium under near-ambient conditions (tube feet, spine and skin) or in small (5000 litre) closed aquarium in artificial seawater (coelomocytes), or (iii) type of RNA-Seq method – stranded RNA-Seq vs CEL-Seq2. These wild-captive differences are more than four times greater than gene expression differences observed between seasons (unpublished) and 173 times greater than between sexes (Jönsson et al., in review). Only metamorphosis has a similar transcriptional change in echinoderms (Wygoda et al., 2014). The scale of transcriptional changes induced by being removed from their natural habitat, and transported to and maintained in captivity likely reflects substantial changes in the physiology, behaviour and health of the COTS. Changes in gene expression in other animals accurately reflects changes in their physiological and developmental in a wide range of contexts (Lv et al., 2013; Meng et al., 2013; Milan et al., 2013; Hui et al., 2014; Gleason and Burton, 2015; Oomen and Hutchings, 2017; Zhang et al., 2017; Li et al., 2019; Ma et al., 2019)

This astonishingly large transcriptional response can be partly attributed to the conditions of captivity leading to an abnormal state of stress (Morgan and Tromborg, 2007; Fischer and Romero, 2019) that can be harmful to organismal homeostasis (Boonstra, 2013; Basile et al., 2021; Hamel et al., 2021). The wide range of potential stressors for captive animals include the abiotic environment (sound, light, odours, temperature), restriction of movement due to confinement, changes in diet, and intra- and inter-species interactions (Morgan and Tromborg, 2007; Jandt et al., 2015). Typically, a stress response consists of physiological, hormonal and behavioural reactions that overcome the imbalance created by the stressors to regain homeostasis, and can manifest as changes in immunological state, metabolism, neural and endocrine systems (Morgan and Tromborg, 2007; Hamel et al., 2021). The stress response caused by being in captivity appears to be highly species-specific, (Dickens and Romero, 2013; Fischer and Romero, 2019). Although most studies to date have been restricted to vertebrates, corals used in laboratory experiments have shown physiological differences compared to corals in the field (Georgiou et al., 2015), suggesting captivity also impacts cnidarians.

### 4.2 Genes consistently up- and downregulated suggest that captive COTS are experiencing oxidative stress

In COTS, 195 CDS are consistently differentially expressed in tube feet, spine, skin and coelomocytes. The 68 upregulated genes appear to be involved in the selective degradation of macromolecules and generation of metabolic energy (catabolism and the ubiquitin system), as well as DNA replication and repair, and protein processing and degradation. This suggests that COTS experience oxidative stress in captivity, where the production and elimination of reactive oxygen species (ROS) are imbalanced, leading to protein, lipid and DNA damage (Sies, 2000; Barzilai and Yamamoto, 2004). The ubiquitin system plays an essential role in regulating translation and protein synthesis to minimise damages caused by ROS (Pickart, 1999; Dougherty et al., 2020). Consistent with captive COTS having high levels of ROS, ubiA prenyltransferase domain-containing protein 1 (an antioxidant enzyme) is upregulated in all four tissues. Energy balance is critical for the survival of organisms under stress (Sokolova et al., 2012). We find that genes involved in energy-releasing catabolic pathways, including the citric acid cycle, fatty acid metabolism and free amino acids pathways, are consistently upregulated in the four tissues in captive COTS, suggesting an increased energy demand. The upregulation of energy metabolic pathways has been observed in marine organisms under environmental stress (Meng et al., 2013; Shekhar et al., 2013; Hui et al., 2014; Nie et al., 2017; Ma et al., 2019; Li et al., 2021) and in captive mussels (Roznere et al., 2021). Given that energy demand appears to decrease under severe stress in some animals (Sokolova et al., 2012), we suggest the apparent increased demand we observe here may reflect a state of moderate chronic stress in these starfish. Consistent with this observation, COTS can be readily kept in captivity for several months (Smith et al., 2017; Smith et al., 2018; Hue et al., 2020).

Among the 127 genes downregulated in all captive COTS tissues are components of conserved cell signalling pathways, including immune, endocrine, nervous and sensory signalling systems, suggesting a general dampening of these systems in captive starfish. The endocrine system is sensitive to environmental changes, and is commonly affected in captive animals (Meijer and Schwabl, 1989; Calisi and Bentley, 2009), and changes in temperature or salinity affect the marine invertebrate neuroendocrine systems (Chen et al., 2008; Massarsky et al., 2011; Zhou et al., 2011; Zhang et al., 2014; Liu et al., 2018). We suggest that the consistent downregulation of genes involved in endocrine and nervous systems in captive COTS may underlie the apparent inability of the starfish to acclimatise after 30 days.

Unlike other aquatic organisms, where signalling pathways appear to be upregulated in response to stress (Lockwood and Somero, 2011; Nie et al., 2017; Yan et al., 2017; Ma et al., 2019), captive COTS downregulate signalling genes in all tissues, such as eight enzymes and fifteen receptors, including five G protein-coupled receptors and two nuclear receptors, both of which also act as a transcription factor. A freshwater mussel in captivity also downregulates receptors, including one found commonly downregulated in the four COTS tissues (Macrophage mannose receptor 1-like) (Roznere et al., 2021). Conversely and consistent with observations in other animals (Lockwood and Somero, 2011; Nie et al., 2017; Yan et al., 2017; Ma et al., 2019), captive COTS upregulate components of Notch and Hippo signalling pathways. The Hippo pathway maintains cellular homeostasis in other bilaterians by regulating stress responses (Di Cara et al., 2015; Mao et al., 2015), suggesting this pathway might play a role in regulating genes involved in oxidative stress in captive COTS.

Six conserved transcription factors (Pax-6-like, RXR, GLIS3, Elf-1-like, PNR, and ZBED1-like) are also downregulated in all four tissues in captive COTS. These may regulate the activation and repression of genes captive starfish across multiple tissues (German et al., 2013; Zhang et al., 2016; Jetten, 2018; Seifert et al., 2019). Oxidative stress appears to mediate the suppression of *Pax6* gene in vertebrates (Zhang et al., 2016). Additionally, RXR receptor, GLIS3 and Elf-1 have all been shown to be involved in signalling in vertebrates, therefore potentially playing a role in the downregulation of cell-cell communication in captive COTS (German et al., 2013; Jetten, 2018; Seifert et al., 2019).

### 4.3 COTS do not appear to acclimatise to their captive environment

Prior to commencing the captivity feeding experiment, COTS were collected on the northern GBR and held in captivity in flow-through aquaria before being air-freighted to Brisbane. In Brisbane, they were held for a further five days in the closed aquarium system in which the experiments were subsequently performed. In total, these starfish were not fed for ten days from initial collection to commencement of the experiment (designated as experimental day 0). After these initial ten days (i.e. day 0 of experiment), 12% of the genes in the entire COTS genome were differentially expressed in coelomocytes, indicating that transport and holding wild COTS in aquaria have a dramatic effect on gene activity. By the end of the experimental period (day 30), 16% of the genes were differentially expressed in the coelomocytes, indicating that COTS do not revert back to natural transcriptional activity after a month in captivity. These results suggest that shorter acclimatisation periods used for COTS (e.g. 2 days - 3 weeks) (Teruya et al., 2001; Petie et al., 2016a; Petie et al., 2016b; Smith et al., 2017; Korsvig-Nielsen et al., 2019) may be insufficient to obtain natural responses to experimental treatments.

Comparison of day 0 and 30 captive and wild coelomocyte transcriptomes suggests that the captive COTS are in different physiological states at these two time points, with genes upregulated at day 0 involved in immunity and genes upregulated at day 30 involved in transcription, DNA repair and ubiquitination, all processes associated with a stress response. This suggests that at day 0 (10 days post-collection) COTS are experiencing acute stress as observed in other animals (Uren Webster et al., 2018), but this state is not sustained over time. 30 days later (40 days post-collection), COTS appear to have progressed into a chronic state of low-grade stress. This is consistent with our interpretation of the observed apparent increase in energy demand, as discussed above.

### 4.4 Captivity appears to induce changes in immune function in COTS

Although changes in immunity appear to be a common response to captivity in vertebrates, this response varies between species with some species experiencing immune suppression while others hyperactivation (Morgan and Tromborg, 2007; Fischer and Romero, 2019). In aquatic invertebrates, disparate changes in stress-induced immune function also have been reported (Lacoste et al., 2002; Gagnaire et al., 2006; Malagoli et al., 2007; Shekhar et al., 2013; Nie et al., 2017; Li et al., 2018; Sun et al., 2019). In captive COTS, most immunity genes appear to be downregulated in all four tissues. The immune response of coelomocytes changes over time. Genes upregulated at day 0 appear involved in pathways similar to the immune response in sea cucumber coelomocytes (leukocyte transendothelial migration, Toll-like receptor signalling pathway, chemokine signalling pathway and Fc gamma R-mediated phagocytosis) (Wu et al., 2020). However, in COTS this immune response is not sustained over time in captivity. Echinoderm coelomocytes are widely used to study innate immunity (Pancer et al., 1999; Pinsino et al., 2007; Gu et al., 2010; Franco et al., 2011; Wu et al., 2020) and this change in the immune response through time suggests that time in captivity should be accounted for when designing those experiments.

## 5 Conclusions

This study reveals that placing COTS in captivity induces massive changes in gene expression that reflect changes in overall homeostatic balance, including energy metabolism, cell signalling and immunity (Figure 4). The large-scale transcriptional changes occur in diverse tissues (tube feet, spine, skin or coelomocytes) and in different types of captive environments (state-of-the-art flow-through aquarium system under ambient conditions, or closed system with artificial seawater, and a constant temperature and light/dark regime). A month in captivity is not sufficient for COTS to return to their native wild transcriptional state. Instead, these starfish appear to progress into a moderate level of chronic stress, which may be maintained for many months. This appears to be similar to the response of freshwater mussels when in captivity (Roznere et al., 2021), suggesting that aquatic invertebrates in general, including corals (Georgiou et al., 2015), are adversely affected by captivity, as observed in many vertebrates. Our findings lead us to urge caution when extrapolating results from captive aquatic invertebrates to their wild counterparts. Specifically, it is important to consider the timing and duration of captivity when designing and interpreting experiments involving COTS and other marine animals. After all, placing species in captivity is subjecting them to conditions never experienced in their evolutionary history.

**Figure 4.**
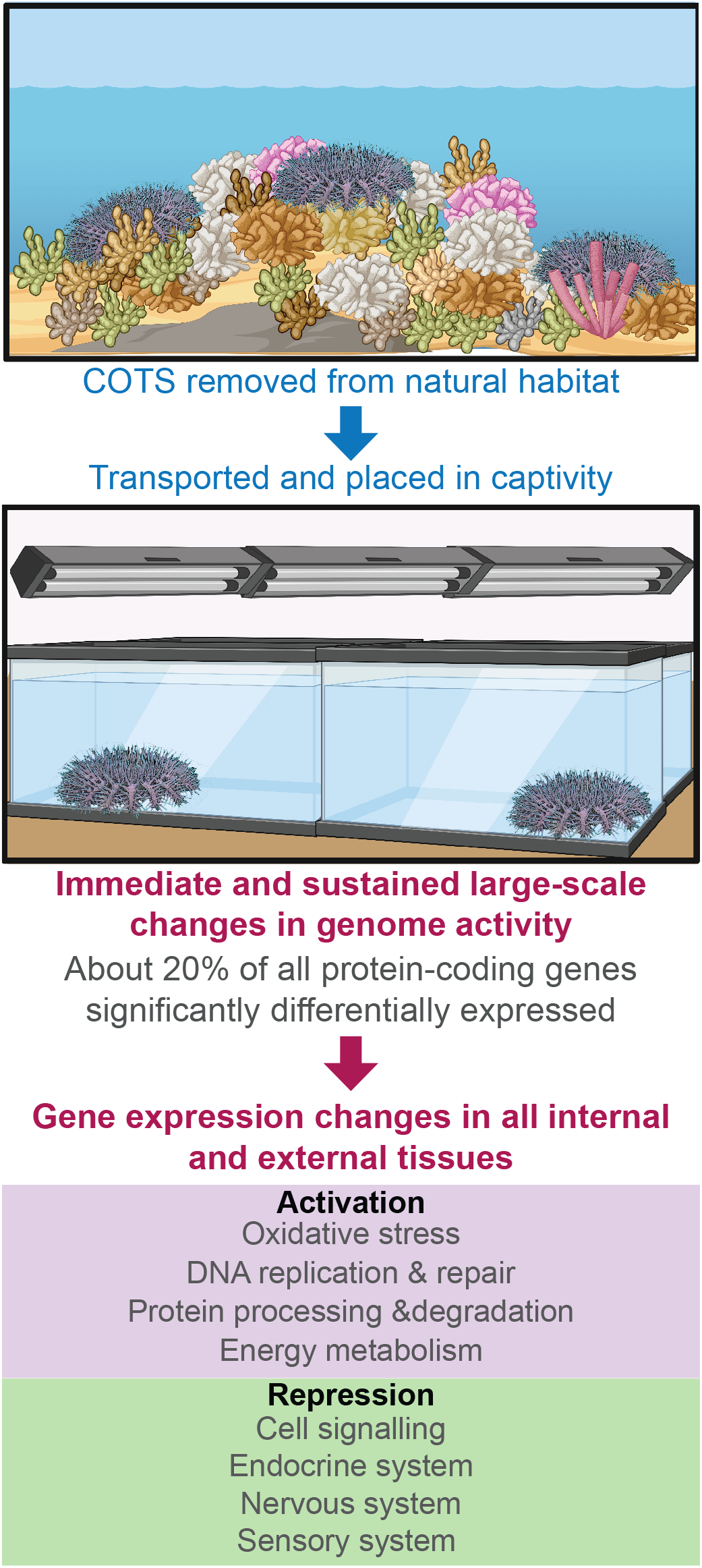
Overview of differential gene expression in crown-of-thorns starfish held in captivity for a month. Transcriptome analyses of crown-of-thorns starfish reveal large-scale changes in global gene expression between wild and captive starfish. Both external and internal tissue types share immediate and sustained large-scale changes in genome activity when transported into captivity, including the activation of genes associated with oxidative stress, replication and repair, protein folding, sorting and degradation, and energy metabolism (purple), and repressing genes associated with cell signalling, endocrine, nervous, and sensory system (green) Taken together, these results highlight the importance of considering the timing and duration of captivity when designing and interpreting experiments involving COTS and other marine animals. This figure was created using Biorender.com.

## 8 Author Contributions

M.M.: Data curation, Formal analysis, Investigation, Visualization, Writing – original draft; M.J.: Data curation, Formal analysis, Investigation, Visualization, Writing – original draft; C.K.W.: Funding acquisition, Writing – review & editing; D.J.C.: Conceptualization, Funding acquisition, Writing – review & editing; S.M.D.: Conceptualization, Funding acquisition, Project administration, Resources, Supervision, Writing – review & editing; B.M.D.: Conceptualization, Funding acquisition, Project administration, Resources, Supervision, Writing – review & editing

All authors gave final approval for publication and agreed to be held accountable for the work performed therein.

## 10 Acknowledgements

We thank AMPTO for collecting COTS and assistance during field trips, Chris Challen for maintaining COTS, and Nick Rhodes for computing assistance.

## 11 Data Availability Statement

The raw RNA sequences generated in this study have been deposited in NCBI Sequence Read Archive (SRA) under BioProject PRJNA821257.

